# Viability of HepG2 and MCF-7 Cells is not Correlated with Mitochondrial Bioenergetics

**DOI:** 10.1101/2022.12.01.518661

**Authors:** Judit Doczi, Noemi Karnok, David Bui, Victoria Azarov, Gergely Pallag, Sara Nazarian, Bence Czumbel, Thomas N Seyfried, Christos Chinopoulos

## Abstract

Alterations in metabolism is a hallmark of cancer. It is unclear, however, if oxidative phosphorylation (OXPHOS) is required for tumor cell survival. We investigated the effect of severe hypoxia, site-specific inhibition of respiratory chain (RC) components, and uncouplers on the survival of HepG2 and MCF-7 2D cultured cells. Comparable respiratory complex activities were observed in both cell lines, but HepG2 cells exhibited much higher oxygen consumption rates (OCR) and respiratory capacity than the MCF-7 cells. Significant non-mitochondrial OCR was found in MCF-7 cells that was insensitive to acute combined inhibition of complexes I and III. However, pre-treatment of either cell line with RC inhibitors for 24-72 hours abolished respective complex activities and OCRs completely, and this was associated with a time-dependent decrease in citrate synthase activity, suggesting mitophagy. HepG2 cells viability was mostly unaffected by any pharmacological treatment or severe hypoxia as temporally recorded from high-content automated microscopy. Conversely, MCF-7 cells viability exhibited strong sensitivity to CIV or CV inhibition, severe hypoxia, and uncoupling, but were only moderately affected by CI, CII and CIII inhibition. CII, CIII and CIV-inhibitor mediated MCF-7 cell death were partially abrogated by aspartate. The data show that OXPHOS activity and viability are uncorrelated in these cell lines indicating that a linkage of OXPHOS to cancer cell survival must be cell- and condition-defined.

## Introduction

Metabolic reprograming for acquiring nutrients to satisfy bioenergetic, biosynthetic, and redox demands is a hallmark of cancer ^1^,^2^. This reprograming is at least partly due to an adverse tumor microenvironment characterized by hypoperfusion and thus, limited [O_2_] partial pressure ^3^. This local hypoxia has fuelled the notion that oxidative phosphorylation (OXPHOS) of tumor cells may be dysfunctional ^4^ the severity of which depend on the actual level of oxygen concentration ^5^; to this end, however, research directives in the past two decades support opposite claims, namely that OXPHOS is either entirely defective ^6^ and this assists growth and metastasis ^7, 8^ or that cancer cells exhibit normal or even enhanced OXPHOS capacity ^9, 10^. More refined statements addressing the role(s) of individual OXPHOS components -such as that of complex I-in cancer cell survival also suffer from the same antipodicity ^11^.

Here we used two of the most widely used cancer cell lines, HepG2 and MCF-7, each cited in >250,000 entries (https://bit.ly/3S5FWee and https://bit.ly/3LfsvWF). Regarding HepG2, originally thought to be a cell culture model of hepatocellular carcinoma, it is now known that these cells rather mimic hepatoblastomas ^12^. Nonetheless, it is a popular hepatic cell line that does not exhibit sublineality ^12^. Furthermore, HepG2 cultures do not exhibit great metabolic variability, thus they are deemed as suitable for real-time assessment of mitochondrial toxicity ^13^.

On the other hand, MCF-7 cells exhibit strong sublineality maintained by the presence of small amounts of estrogen in the fetal bovine serum, thus being at the mercy of the vendor and batch-to-batch variability ^14^. In addition to that, MCF-7 cells sublines exhibit differences in copy number alteration profiles, affecting even the most basic aspects of cell phenotype ^15^. Nonetheless, most laboratories -except the group of Lisanti ^16, 17, 18^, and another group in China ^19^-concur on the findings that MCF-7 cells respire minimally ^20, 21, 22, 23, 24, 25, 26^.

We report that HepG2 cells exhibit robust OCRs and respiratory capacity, implying OXPHOS-capable *in situ* mitochondria; yet, inhibition of any of the respiratory complexes leading to a complete loss of respiration, or severe hypoxia or uncoupler-conferred complete collapse in ΔΨm exerted no effect on their viability. On the other hand, MCF-7 cells exhibited very little respiratory capacity implying OXPHOS-defective *in situ* mitochondria; however, some RC inhibitors, severe hypoxia or maximal uncoupling led to loss of MCF-7 cells viability. Furthermore, MCF-7 cells also showed a non-mitochondrial component consuming oxygen. As no pattern emerged, thus no correlation between OXPHOS capacity and cell viability could be established, we posit that the reliance of tumor cells to OXPHOS for survival is cell- and condition-dependent. We further caution that regarding the relation of mitochondrial oxidative phosphorylation and survival no safe extrapolations can be made from one cancer cell type to another.

## Materials and Methods

### Cell cultures

HepG2- and MCF7 cells were grown in Dulbecco’s Modified Eagle Media and in Minimum Essential Medium Eagle, respectively. All cultures were supplemented with 10% fetal bovine serum, 2 mM glutamine and kept at 37 °C in 5% CO_2_. All media were further supplemented with penicillin, streptomycin and amphotericin. Experiments were carried out at day 1–2 post plating, unless stated otherwise.

### Isolation of mitochondria from cell cultures

Cells were harvested by scraping and centrifuged (2 min, 3,500 rpm, 4°C); pellets were saved, supernatants were re-spun and pellets were combined. The pellet was resuspended in a solution containing 225 mM mannitol, 75 mM sucrose, 5 mM HEPES, 1 mM EGTA, 1 g/l bovine serum albumin, pH 7.4 centrifuged again and the supernatant was discarded. The pellet was resuspended in the same buffer and homogenised with 10 strokes of A and 5 strokes of B pestle in a Dounce homogeniser. The homogenate was centrifuged twice (10 min, 600g, 4°C) and the pellet was discarded. Afterwards the supernatant was centrifuged (10 min, 14,000g, 4°C). The supernatant was discarded and the pellet was resuspended in the original media that was saved.

### OXPHOS complex assays

10^5^ cells were seeded in 25 cm^2^ flasks (for complexes II, III, IV) and after one day (or upon reaching 90% confluence for complex V) the culture medium was exchanged with a solution the composition of which was: 120 mM NaCl, 3.5 mM KCl, 20 mM HEPES, 1.3 mM CaCl_2_, 1 mM MgCl_2_, 10 mM glucose, 4 mM glutamine, pH 7.4 with or without OXPHOS inhibitor (5 μM rotenone, 1 μM atpenin A5, 1 μM myxothiazol 1 mM NaN_3_, or 5 μM oligomycin). For CI measurement cells were seeded in 175 cm^2^ flasks and grown to near confluence before medium change. After 24h, 48h or 72h of incubation, cells were harvested by scraping, centrifuged (2 min, 3500 rpm, 4°C) and resuspended in a lower volume of their respective supernatant except for CI measurement, where isolated mitochondria were prepared. Samples were permeabilized by 3 freeze-thaw cycles. Complex I (CI), Complex II (CII), Complex III (CIII) and Complex IV (CIV) assays were performed as described in ^27^ and ^28^ with the recommendations outlined in ^29^; concentrations were adjusted to suit our instrumentation (Tecan Infinite® 200 PRO series plate reader, Tecan Deutschland GmbH, Crailsheim, Germany) using 96 well plates. Transparent flat bottom plates were used for absorbance measurements.

### CI (NADH:decylubiquinone oxidoreductase)

Permeabilized mitochondria were added to two reaction mixtures (25 mM KH_2_PO_4_, 2.5 g/l BSA, 0.3 mM NaCN, 0.1 μM myxothiazol, pH 7.2) with or without 1 μM rotenone. The final volume is double that of the sample added. After 10 min of temperature equilibration at 37°C, 120 μM decylubiquinone was added. Reaction was started with 100 μM NADH. Initial NADH oxidation was measured at 340 nm and reaction rate was calculated as the rotenone-sensitive NADH oxidation (*ε*_NADH_=6.2 mM^-1^·cm^-1^*l*=0.7 cm) at the highest signal-to-noise ratio interval. CII (succinate:2,6-dichlorophenolindophenol oxidoreductase): The permeabilized cell suspension was added to two reaction mixtures (25 mM KH_2_PO_4_, 20 mM succinate, 0.3 mM NaCN, 0.1 μM myxothiazol, 1 μM rotenone, pH 7.2) with or without 10 mM malonate. The final volume is double that of the sample added. After 10 min, 65 μM 2,6-dichlorophenolindolphenol (DCPIP) was added. Reaction was started with 50 μM decylubiquinone. Initial DCPIP reduction was measured at 600 nm and reaction rate was calculated as the malonate-sensitive DCPIP reduction (*ε*_DCPIP_=19.1 mM^-1^·cm^-1^ *l*=0.7 cm).

CIII (decylubiquinol:ferricytochrome C oxidoreductase) activity was assayed similar to CII with a reaction mixture (50 mM KH_2_PO_4_, 1 g/l BSA, 0.3 mM NaCN, 1 μM rotenone, 0.1 mM EDTA, 1 mM n-dodecyl-β-D-maltoside, pH 7.2) with or without 0.1 μM myxothiazol. Without delay, 92.5 μM decylubiquinol was added. Reaction was started with 50 μM ferricytochrome C. Initial ferricytochrome C reduction was recorded at 550 nm and reaction rate was calculated as the myxothiazol-sensitive cytochrome c reduction (Δ*ε*_cytc_=18.5 mM^-1^·cm^-1^ *l*=0.7cm).

CIV (ferrocytochrome c oxidase) activity was assayed similar to CII with a reaction mixture (20 mM KH_2_PO_4_, 0.1 μM myxothiazol, 0.45 mM n-dodecyl-β-D-maltoside, pH 7.0) and with or without 5 mM azide. After 10 min of temperature equilibration on 37°C, reaction was started with 50 μM ferrocytochrome C. Cytochrome C was reduced freshly on the day of the measurement with dithionite. Initial cytochrome C oxidation was recorded at 550 nm. After the measurement concentrated ferricyanide was added and the absorption was recorded. Azide-sensitive pseudo first order rate constant was calculated from the ferrocytochrome c oxidation.

### CV (F_o_-F_1_ ATPase)

The permeabilized cell suspension was added to two reaction mixtures (50 mM Tris, 2 mM EGTA, 1 mM MgCl_2_, pH 8.0) with or without 50 μM oligomycin. After 5 min of temperature equilibration at 37°C measurement was started while constant shaking. Reaction was started with 2 mM ATP. Samples were taken at 1, 2 and 3 minutes and the reaction was quenched with equal volume of 10 w/v% trichloroacetic acid. After centrifugation (5 min, 14,000 rpm) 25 μl supernatant was added to 155 μl colour reagent (45 parts of 13.33 mM molybdenum(VI), 0.44 mM antimony(III) in 1.33 mM tartarate; 110 parts of dimethyl sulfoxide) parallel to phosphate standards. Colour development was started by adding 20 μl 1 w/v% ascorbic acid and after 15 min the absorbance was read at 890 nm. Reaction rate was calculated as the oligomycin-sensitive ATP hydrolysis to phosphate.

### Citrate synthase

The sample was added to a reaction mixture (100 mM Tris, 0.1 mM 5,5′-Dithiobis(2-nitrobenzoic acid, DTNB), 0.1 w/v% Triton-X100, pH 8.0). After 10 min of temperature equilibration at 37°C, 0.1 mM acetyl-CoA was added. Reaction was started with 0.25 mM oxaloacetate. Initial DTNB reduction was recorded at 412 nm and reaction rate was calculated directly; background thiolase activity was in all cases negligible (*ε*_DTNB_ =13.6 mM^-1^·cm^-1^ *l*=0.7cm)

Oxygen consumption and extracellular acidification rates in cell cultures: Real-time measurements of oxygen consumption rate (OCR) and extracellular acidification rate (ECAR) were performed on a microfluorimetric XF96 Analyzer (Seahorse Bioscience, North Billerica, MA, USA) as previously described ^30^. Cells were seeded 1 day before addition of the inhibitors (or vehicle) in Seahorse XF96 cell culture microplates at ∼2*10^4^ cells/well density, in growth media. At 0 hours growth media were changed and the inhibitors (or vehicle) were added in assay media containing (in mM): 120 NaCl, 3.5 KCl, 1.3 CaCl_2_, 1.0 MgCl_2_, 20 HEPES, 10 glucose, 4 glutamine, pH 7.0 and the experiments were conducted at 24, 48 and 72 hours. OCR and ECAR values were calculated by the XF96 Analyzer software. During the measurement, 20-26 μl of testing agents prepared in assay media were then injected into each well to reach the desired final working concentration.

### Cell viability and total cell number

HepG2 or MCF-7 cultures were cultured in poly-D-Lysine coated Corning 96-well High Content Imaging Glass Bottom Microplate (#No. 4580, Corning Inc., USA) for 1-2 days, at a density of approximately 2*10^4^ cells/well. Cultures were incubated in a standard CO_2_ incubator at 37°C in assay medium containing (in mM): 120 NaCl, 3.5 KCl, 1.3 CaCl_2_, 1.0 MgCl_2_, 20 HEPES, 10 glucose, 4 glutamine at pH 7.4 and moved into the microscope for further imaging at room temperature and atmospheric CO_2_ concentration at the indicated time-points. For representative images and nucleus-based quantification of live and dead cells, a nuclear staining with 1 μg/ml Hoechst 33342 dye (Invitrogen,Thermo Fisher Scientific, Corp., USA) and propidium iodide dye (0.1 μg/ml, Invitrogen) were added to assay media for 20 min before the image acquisition. Cell cultures were imaged with an ImageXpress Micro Confocal High Content Imaging System (Molecular Devices). Filter pairs of 377/50 and 447/60 nm for Hoechst 33342 and 562/40 and 624/40 nm were used for propidium iodide. Imaging of one 96-well plate took approximately 40 min which the cells seemed to tolerate reasonably well. During image acquisition, nine 20× magnified fields of view (each covering 0.7209 mm^2^) were captured from each well and at least four parallel wells were imaged for each condition. Images were quantified with MetaXpress High Content Image Acquisition & Analysis Software to determine the number of objects of interest based on intensity above local background as well as minimal and maximal object sizes. The cell scoring workflow used attempted to separate touching/overlapping suprathreshold objects as well, thus the total object count was used for further quantitative characterization of the cultures. Detection and quantification of nuclei was likewise performed in the MetaXpress software using a built-in algorithm that determined the total number of nuclei identified by scoring workflow of the blue and red channel images.

Simultaneous measurement of mitochondrial (ΔΨm) and plasma membrane potential (PMP): HepG2 or MCF-7 cultures were grown for imaging on poly-D-Lysine coated 8-well LabTek II chambered coverglasses (Nunc, Rochester, NY, USA) for 1-2 days, at a density of approximately 3*10^4^ cells/well. Cultures were incubated at 37°C in imaging medium containing (in mM): 120 NaCl, 3.5 KCl, 1.3 CaCl_2_, 1.0 MgCl_2_, 20 HEPES, 10 glucose, 4 glutamine, pH 7.4 with TMRM (180 nM) plus the bis-oxonol type plasma membrane potential indicator, DIBAC_4_(3) (250 nM, Life Technologies Inc.) for 60 min before the experiment. Experiments were performed at 34 °C on an Olympus IX81 inverted microscope equipped with a UAPO 20 × 0.75 NA lens, a Bioprecision-2 xy-stage (Ludl Electronic Products Ltd., Hawthorne, NY) and a 75W xenon arc lamp (Lambda LS, Sutter Instruments, Novato, CA). Time lapses of 1342 × 1024 pixels frames (digitized at 12 bit with 4 × 4 binning, 250 msec exposure time for TMRM and 100 msec exposure time for DIBAC_4_(3) were acquired by an ORCA-ER2 cooled digital CCD camera (Hamamatsu Photonics, Hamamatsu, Japan) under control of MetaMorph 6.0 software (Molecular Devices; Sunnyvale, CA, USA). For the illumination of DIBAC_4_(3) a 490/10 nm exciter, a 505LP dichroic mirror and 535/25 emission filter were used, for TMRM a 535/20 nm exciter, a 555LP dichroic mirror and a 630/75 emitter were used, all from Chroma Technology Corp., (Bellows Falls, VT, USA). The cross-talk of TMRM and DIBAC_4_(3) emissions was eliminated by a linear spectral un-mixing algorithm implemented in Image Analyst MKII (Image Analyst Software, Novato, CA) as previously described ^31^, using decomposition algorithms developed in ^32^. The algorithm calculates un-mixed fluorescence intensities by solving a linear equation with the cross-talk coefficient matrix. Coefficient matrix was determined by loading the cultures either with TMRM or DIBAC_4_(3) while both emissions (TMRM and DIBAC_4_(3)) were recorded. At the end of each experiment, full mitochondrial depolarization and plasma membrane depolarization was achieved by the application of mitochondrial (MDC) and plasma membrane depolarization cocktails (CDC). MDC contained (in μM): 1 valinomycin, 1 SF 6847, 2 oligomycin; CDC contained (in μM): 1 valinomycin, 1 SF 6847, 2 oligomycin, 10 nigericin, 10 monensin.

Hypoxia treatment of cells: Plated cells were placed in the interior of a hypoxia incubator chamber (STEMCELL technologies) and flushed with 95% Nitrogen (purity: 99,99999%) and 5% CO_2_ for 4 min, at 1.4 Bar. Subsequently, the chamber was hermetically closed and placed within a thermocontrolled incubator, at 37 °C. After 40 min, the process was repeated once and the cells were returned to the thermocontrolled incubator. Cells were processed at 24, or 48 or 72 hours after this hypoxic treatment.

### Verification of hypoxia

For evaluating the extent of hypoxia in cells, cultures were treated with EF5 (2-(2-Nitro-1H-imidazol-1-yl)-N-(2,2,3,3,3-pentafluoropropyl) acetamide), a compound that binds to hypoxic cells and forms adducts ^33^. The adducts were detected by a mouse monoclonal antibody (clone ELK3-51) conjugated to Alexa 488. The fluorescence of Alexa 488 was visualized by epifluorescence microscopy.

### Animals

Mice were of mixed 129 Sv and C57BL/6 background. The animals used in our study were of either sex and between 2 and 6 months of age. Data obtained from liver mitochondria of mice of a particular gender or age (2, 4, or 6 months) did not yield any qualitative differences, thus all data were pooled. Mice were housed in a room maintained at 20–22 °C on a 12-h light–dark cycle with food and water available *ad libitum*. The study was approved by the Animal Care and Use Committee of the Semmelweis University (Egyetemi Állatkísérleti Bizottság, protocol code F16-00177 [A5753-01]; date of approval: May 15, 2017).

### Isolation of mouse liver mitochondria

Liver mitochondria were isolated from mice as described in ^34^.

Determination of membrane potential (ΔΨm) in isolated mitochondria: ΔΨm of isolated mitochondria (1 mg of mouse liver mitochondria in two ml buffer medium, the composition of which is described in ^35^) was estimated fluorimetrically with 5 μM rhodamine 123^35^. Fluorescence was recorded using an Oroboros O2k (Oroboros Instruments, Innsbruck, Austria) equipped with the O2k-Fluo LED2-Module with optical sensors including an LED (465 nm; <505 nm short-pass excitation filter), a photodiode, and specific optical filters (>560 nm long-pass emission filter) ^36^. The experiments were performed at 37 °C.

### Determination of respiration in isolated mitochondria

Oxygen consumption was performed polarographically using an Oroboros O2k simultaneous to determination of ΔΨm, thus conditions were identical. The oxygen concentration (μM) and oxygen flux (pmol·s^-1^·mg^-1^; negative time derivative of oxygen concentration, divided by mitochondrial mass per volume and corrected for instrumental background oxygen flux arising from oxygen consumption of the oxygen sensor and back-diffusion into the chamber) were recorded using DatLab software (Versions 7.3.0.3 Oroboros Instruments, Innsbruck, Austria).

### Protein determination

The protein equivalent concentration was determined using the bicinchoninic acid assay and calibrated using bovine serum standards ^37^ using a Tecan Infinite® 200 PRO series plate reader (Tecan Deutschland GmbH, Crailsheim, Germany).

### Reagents

Standard laboratory chemicals and digitonin were from Sigma. SF 6847 was from Biomol (BIOMOL GmbH, Hamburg, Germany). Fetal bovine serum was from Atlanta Biologicals (Lawrenceville, GA, USA) and all other tissue culture reagents were purchased from Life technologies.

### Statistics

Data are presented as mean ± SEM or ± SD as stated in the legend; significant differences between two groups of data were evaluated by Student’s t-test, with p < 0.05 considered significant. Significant differences between three or more groups of data were evaluated by one-way analysis of variance followed by Tukey’s post-hoc analysis, with p < 0.05 considered statistically significant. If normality test failed, ANOVA on Ranks was performed.

## Results

### Respiratory chain inhibitors confer a complete and persistent inhibition of their targets throughout the experimental time frame

A main aim of the present study is to investigate the effects of site-specific RC inhibition on HepG2 and MCF-7 cells viability in a temporal manner. For this, it must be ensured that RC inhibition by the drugs is complete and sustained throughout the experiments, *i*.*e*. respiratory complex inhibition is not overcome by cellular proliferation, biotransformation and/or decomposition of the inhibitor(s) within the 72 hour time frame. As shown in figure 1 for HepG2 cells and figure 2 for MCF-7 cells, CI, CII, CIII and CIV activities normalized to protein (A, C, E, G) or citrate synthase (CS) activity (B, D, F, H) the inhibitors (rotenone for CI, atpenin A5 for CII, myxothiazol for CIII and azide for CIV) sustained complete inhibition of the respective complexes for the entire duration of the experiment (grey bars), compared to untreated cells (black bars). An exception to this statement is the effect of CII inhibitor atpenin A5 on MCF-7 cells after incubation for 3 days (figure panels 2C, D); there, CII activities seem to rebound by 20-30% after experiencing complete abolition in the first two days. Due to the crudeness of the homogenates the specificity of the activity assay was limited to around 40-50% for CI and 60-70% for CV. In addition, the decrease in protein equivalent or citrate synthase (CS) activity over time tended to magnify the variance and error. From the data shown in figure panels 1A-H and 2A-H it is also evident that the activities of most complexes (except CIII in MCF-7 cells) decreases over time, in the absence of the inhibitors. This latter phenomenon was not investigated further.

**Figure 1.**
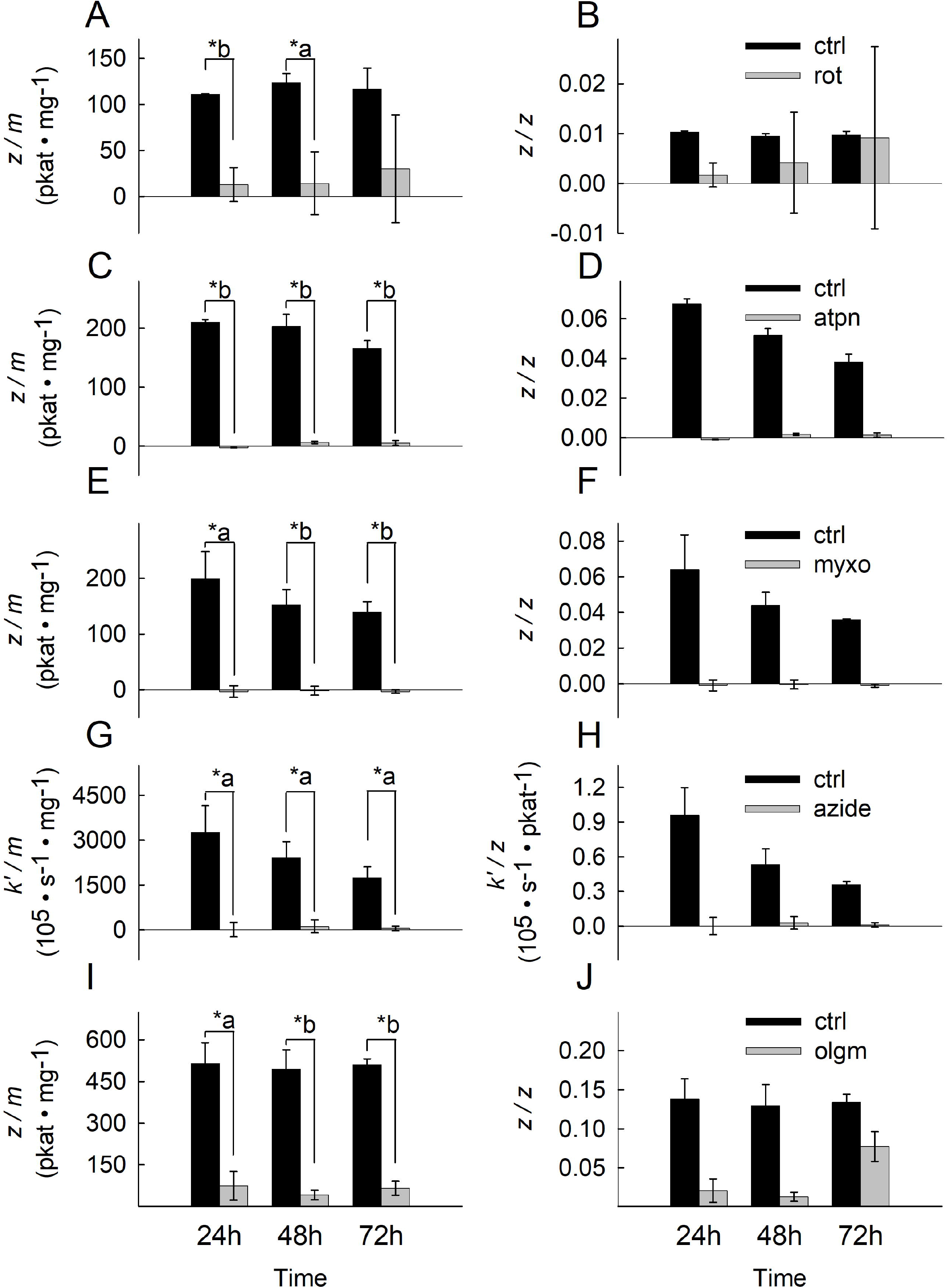
Effect of site-specific inhibitors on OXPHOS component activities in HepG2 cells over time. Bars depicting CI, CII, CIII, CIV and CV inhibitor-sensitive catalytic activities normalized to protein equivalent (pkat / mg) (**A, C, E, G, I**) or citrate synthase (CS) activity (unitless) (**B, D, F, H, J**) after incubation for 24h, 48h and 72h at 37 °C under 5% CO_2_ atmosphere in assay media containing the respective RC inhibitor. The inhibitors (5 μM rotenone for CI, 1 μM atpenin A5 for CII, 1 μM myxothiazol for CIII, 1 mM azide for CIV, 5 μM oligomycin for CV) sustained complete or near complete inhibition of the respective complex for the entire duration of the experiment (grey bars), compared to untreated cells (black bars). Data indicate the mean of three biological replicates and error bars indicate 1 standard deviation. *a: p <0.05; *b: p <0.001.

**Figure 2.**
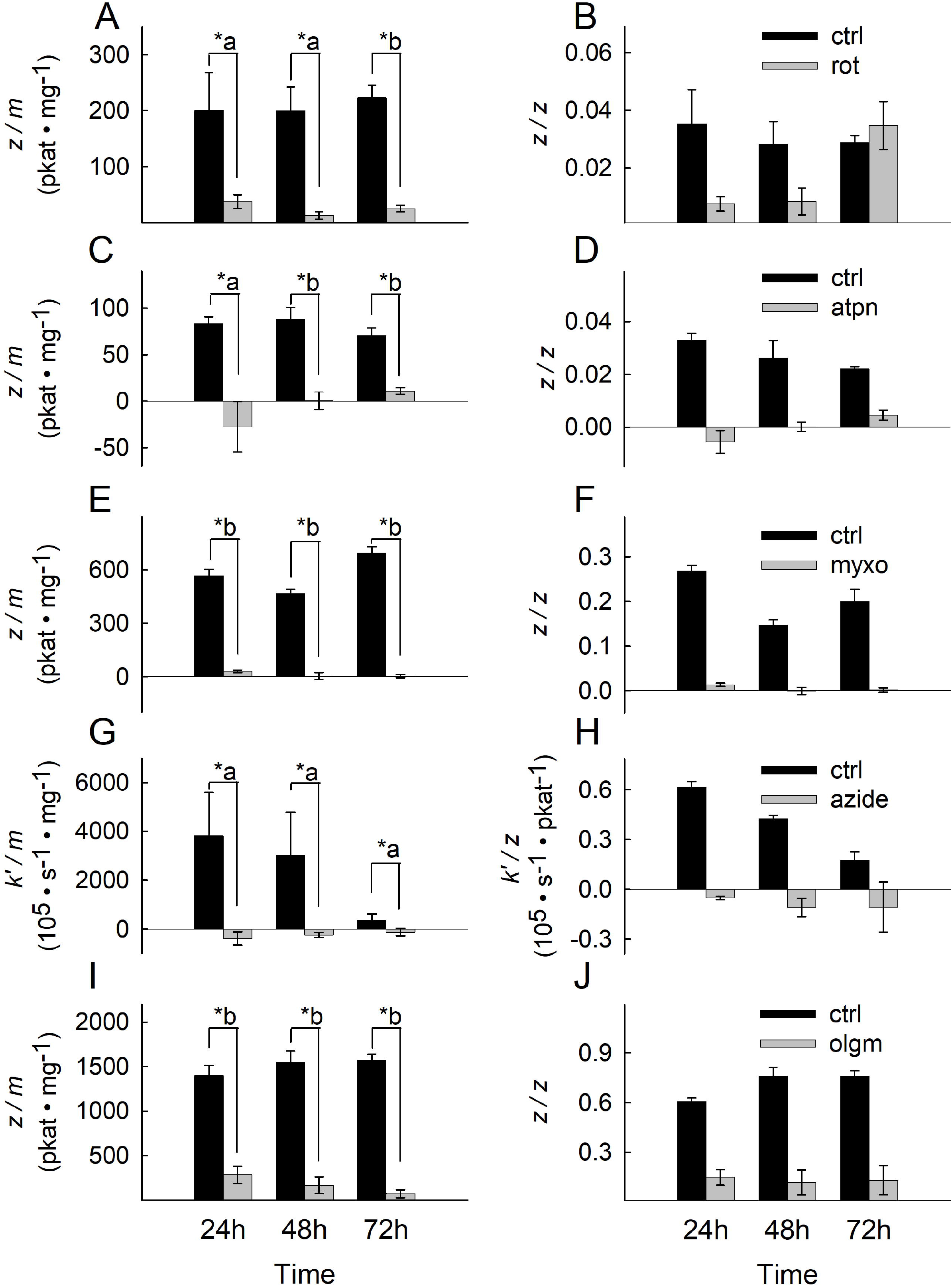
Effect of site-specific inhibitors on OXPHOS component activities in MCF7 cells over time. Bars depicting CI, CII, CIII, CIV and CV inhibitor-sensitive catalytic activities normalized to protein equivalent (pkat / mg) (**A, C, E, G, I**) or citrate synthase (CS) activity (unitless) (**B, D, F, H, J**) after incubation for 24h, 48h and 72h at 37 °C under 5% CO_2_ atmosphere in assay media containing the respective RC inhibitor. The inhibitors (5 μM rotenone for CI, 1 μM atpenin A5 for CII, 1 μM myxothiazol for CIII, 1 mM azide for CIV, 5 μM oligomycin for CV) sustained complete or near complete inhibition of the respective complex for the entire duration of the experiment (grey bars), compared to untreated cells (black bars). Data indicate the mean of three biological replicates and error bars indicate 1 standard deviation. *a: p <0.05; *b: p <0.001.

Of note, we added azide instead of the most commonly used cyanide for inhibiting CIV. The reason for this is because we discovered that on neutral pH cyanide evaporates significantly over the course of several hours, thus its potency in inhibiting CIV is decreasing, at least within the experimental time frame. Azide did not suffer from the same limitation; on the other hand, we also discovered that azide exhibits uncoupling properties in higher doses, as shown in supplementary figure 1. So we opted for a 1 mM concentration as a compromise between uncoupling and inhibition.

The activity of CV (F_o_-F_1_ ATPase) is given as the oligomycin-sensitive ATP hydrolysis rate; as shown in figure panels 1I, J (for HepG2 cells) and figure panels 2I, J (for MCF-7 cells) CV activity normalized to protein or CS activity is evident in both cell lines. From the above data we conclude that both cell lines exhibit CI-CV activities and that the CI-CV inhibitors sustain inhibition of the intended complex throughout the experimental time frame.

RC inhibitors maintain abolition of oxygen consumption rates within the experimental time frame Having established that all drugs sustain inhibition of the respective complexes within the experimental time frame, we sought to establish the effect of complexes inhibition on *in situ* mitochondrial bioenergetic competence. For this, we first evaluated the effects of the inhibitors on oxygen consumption rates (OCR) and extracellular acidification rates (ECAR) in intact, adhering cells in 2D cultures. The inhibitors were added at 0 hours and the experiments were conducted at 24, 48 and 72 hours. Regarding CI, we used three different inhibitors: rotenone (rot), piericidin A (pierc) and pyridaben (prdbn). The reason for this is because rotenone has been described to inhibit microtubule assembly ^38, 39, 40, 41, 42^, which in turn may exert an effect of cell adherence ^43^; if cells detach from the plates that would lead to the result of showing less oxygen consumption from the vicinity of the plate bottom where adherent cells are expected to reside.

As shown in figure panels 3A, C, E for HepG2 cells, each and every RC inhibitor included in the media for the entire duration of the experiments i) diminished basal OCR, ii) abolished the oligomycin (olgm)-induced decrease in OCR and iii) abolished the uncoupler (by DNP)-induced increase in OCR. Thus, all RC inhibitors prevented completely intact *in situ* mitochondria from consuming oxygen. Because each and every RC inhibitor abolished OCR, this means that OCR was almost exclusively mitochondrial; the validity of this statement is only limited by the detection threshold of OCR, i.e. the Seahorse 96 analyzer. This is also in line with the finding that addition of antimycin and rotenone (A+R) shown in the end of every Seahorse experiment decreased OCR to very low (but not zero) levels. However, it is surprising that inhibition of CII also led to complete abolition of OCR; one may have expected that CII blockade would have let the CI -> CIII -> CIV electron flow unaffected, but this was not the case. On the other hand, as expected, combined inhibition of CI+ CV or CIII+CV led to nearly complete collapse of ΔΨm, shown in supplementary figure 2. In line with the OCR data, ECAR experiments performed in the same HepG2 cells at the same time shown in figure panels 3B, D, F at 24, 48 and 72 hours respectively demonstrate that i) each and every RC inhibitor present in the media led to an increase in ECAR, and ii) where no RC inhibitor was present, addition of oligomycin led to a rebound increase in ECAR, reaching levels as with the 24-72 hour long treatment with RC inhibitors. The results support the conclusion that RC inhibitors abolished OXPHOS that led to compensatory increases in glycolytic fluxes yielding pyruvate and lactate formation, acidifying the media. The lack of effect of adding glucose and glutamine (indicated in the panels) on OCR and ECAR attests to the fact that cells were not starved by either substrate.

OCR of MCF-7 cells was much less than that of HepG2, in accordance to most other reports (see Introduction). Modest oligomycin-induced decreases followed by DNP-induced increases in OCR (in the absence of other RC inhibitors) is observed in figure panels 4A, C and E (black symbols). The presence of each and every RC inhibitor abolished all these responses. The results support the conclusion that our MCF-7 sub-lineal cultures exhibit poor OCR indicating very weak OXPHOS. Furthermore, it is noteworthy that addition of A+R to MCF-7 cells did not yield near zero OCR levels, thus, this pre-A+R oxygen consumption cannot be attributed to mitochondria. Pre-treatment of cultures for 24-72 hours with any RC inhibitor however, resulted in the loss of the basal OCR rates. The latter findings attest to a serious pitfall: OCR of MCF-7 cultures may not be entirely mitochondrial but 24-72 hour treatment with RC inhibitors abolishes the non-mitochondrial origin of oxygen consumption.

The ECAR data obtained from MCF-7 cells (also performed in the same cells at the same time as for OCR) shown in figure panels 4B, D F reveal very low rates in pH alterations in the media for any solid conclusions to be drawn.

From figure 4 it is also evident that 1-3 day pre-treatment with any RC inhibitor yields less OCR and ECAR values compared to the acute addition of A+R. To address this further, we compared citrate synthase activities as means of mitochondrial content as a function of RC inhibitor pre-treatment time. As shown in supplementary figure 3, there is a time-dependent decrease of citrate synthase activity in RC inhibitor treated cells with the exception of atpenin, compared to the un-treated cells. This means that RC inhibitor treatment for 1-3 days diminishes mitochondrial content. We conservatively attribute this to mitophagy ^44^; this phenomenon was not investigated further.

**Figure 3:**
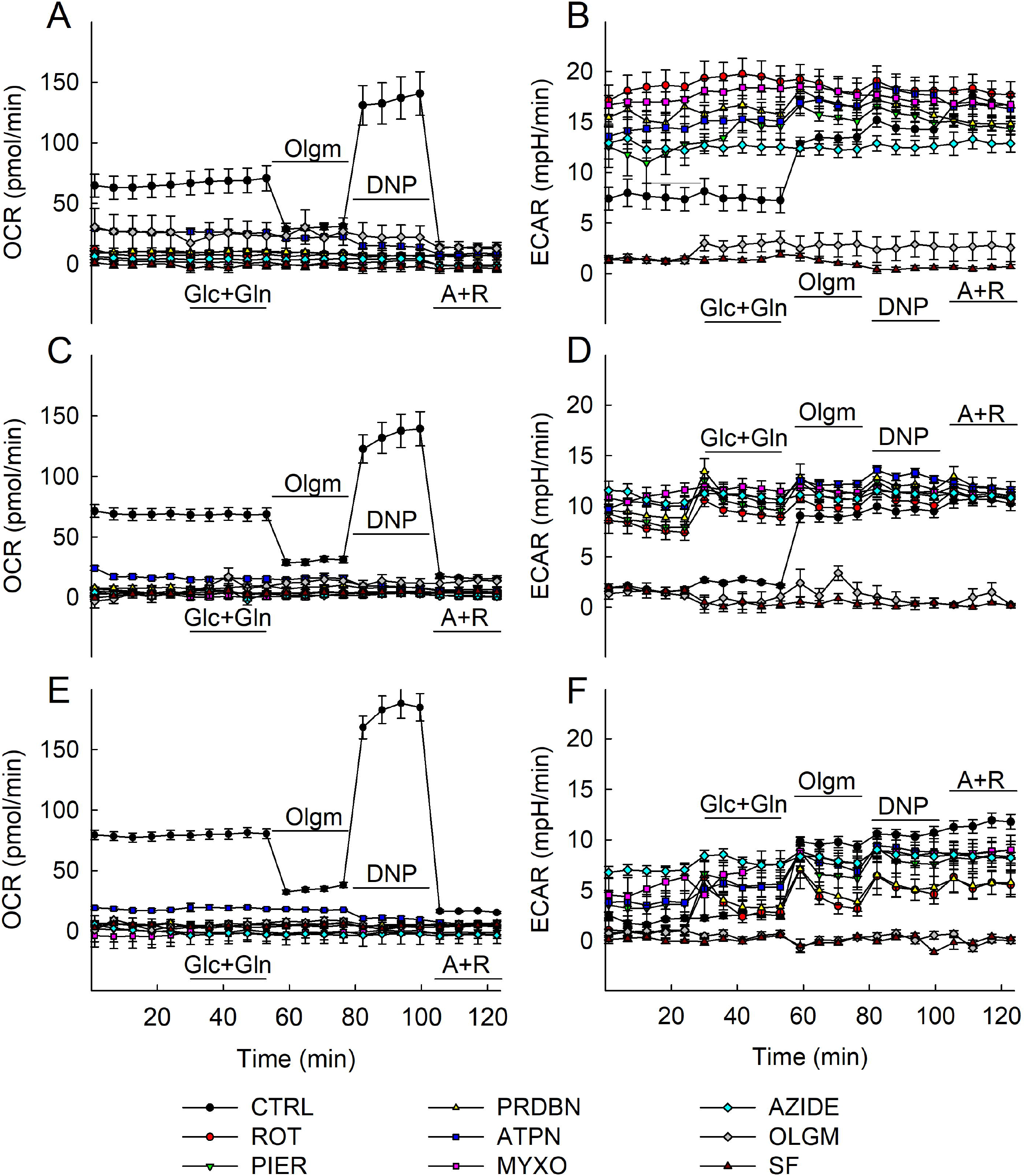
Metabolic profiling and effects of RC inhibitor additions on mitochondrial and glycolytic activities in HepG2 cells. HepG2 cells were cultured in XF96-well cell culture microplates (Seahorse Bioscience) at a density of 2 × 10^4^ cells/per well and then incubated for 24 h (**A**,**B**), 48h (**C**,**D**) and 72h (**E**,**F**) at 37 °C under 5% CO_2_ atmosphere in assay media containing RC inhibitors at the following concentrations: rotenone 5 μM (ROT), piericidin 1 μM (PIER), pyridaben 1 μM (PRDBN), atpenin 1 μM (ATPN), myxothiazol 1 μM (MYXO), azide 1 mM (AZIDE), oligomycin 5 μM (OLIGO) and SF 6847 1 μM (SF). OCR (**A**,**C**,**E**) and ECAR (**B, D**,**F**) values were normalized to total cell numbers in the assay. Data are representative of at least three independent experiments, each additions with 12 statistical replicates and error bars indicate SEM.

**Figure 4:**
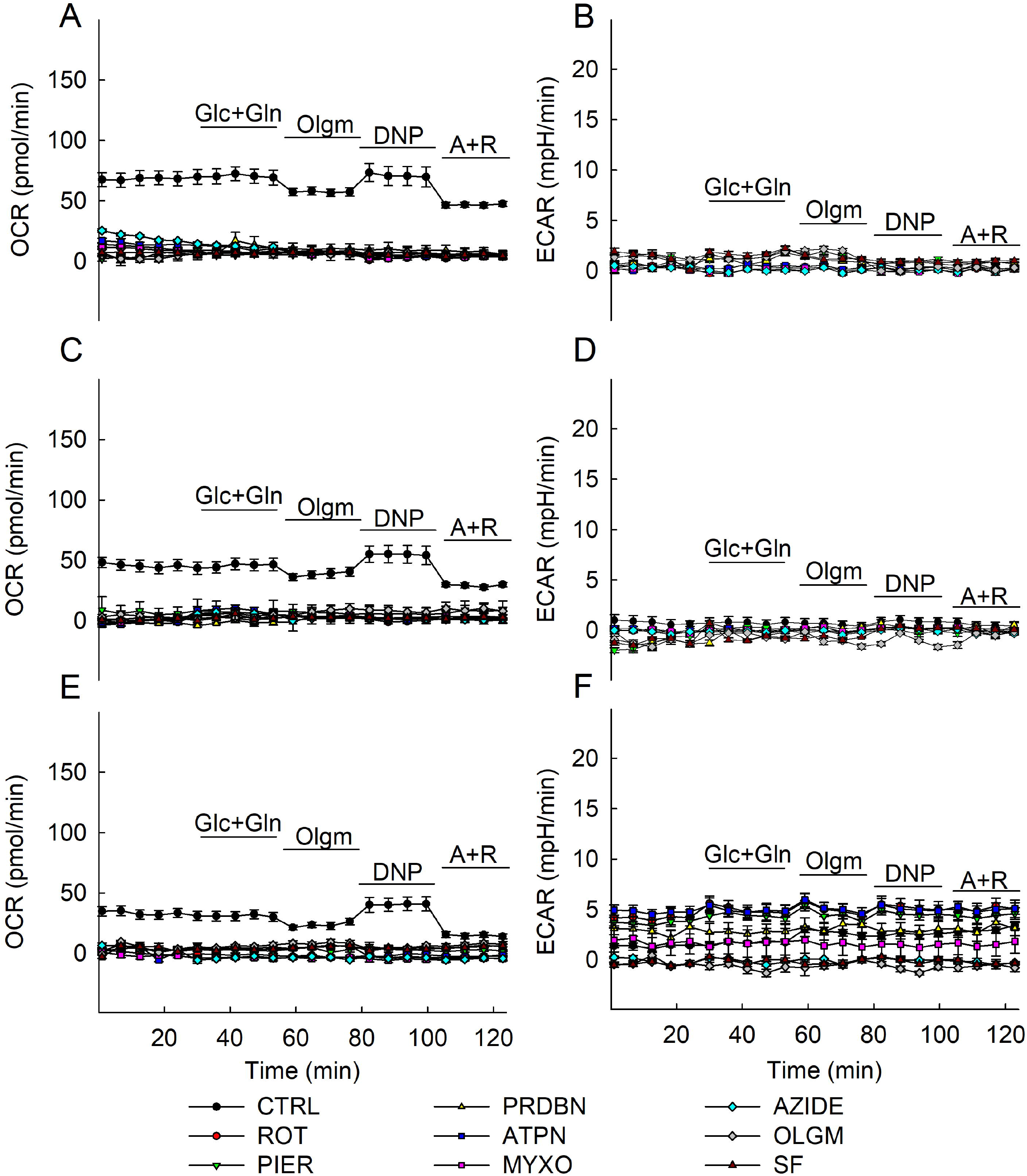
Metabolic profiling and effects of RC inhibitor additions on mitochondrial and glycolytic activities in MCF7 cells. MCF7 cells were cultured in XF96-well cell culture microplates (Seahorse Bioscience) at a density of 2 × 10^4^ cells/per well and then incubated for 24 h (**A**,**B**), 48h (**C**,**D**) and 72h (**E**,**F**) at 37 °C under 5% CO_2_ atmosphere in assay media containing RC inhibitors. Concentrations of RC inhibitors were identical to those indicated in the legend of Fig. 3. OCR (**A**,**C**,**E**) and ECAR (**B, D**,**F**) values were normalized to total cell numbers in the assay. Data are representative of at least three independent experiments, each addition with 12 statistical replicates and error bars indicate SEM.

### Titration of the uncoupler SF6847 as a function of *in situ* mitochondrial membrane potential (ΔΨm)

Prior to testing the effect of maximum uncoupling on cell viability, we titrated the uncoupler SF6847 and recorded TMRM fluorescence (reporting both plasma membrane and mitochondrial membrane potential) subtracted from the DIBAC_4_(3) fluorescence (reporting only plasma membrane potential) in HepG2 and MCF-7 2D cultures using wide-field epifluorescence. As shown in figure panels 5A and 5B for HepG2 and MCF-7 cell cultures respectively, the uncoupler SF6847 dose-dependently decreased the fluorescence values. When this decrease started to exhibit a plateau phase, a membrane potential-dissipating cocktail - MDC, the composition of which is detailed under materials and methods-was applied to ensure that no further loss of fluorescence could be achieved. From the experiments shown in figure panels 5A and 5B we deduced that 1 μM SF6847 was sufficient to confer complete uncoupling in both cell culture types.

### The effect of bioenergetic impairment on HepG2 and MCF-7 cells viability

Having established that the RC inhibitors abolish the activity of respective complexes completely and eliminate OCRs throughout the experimental time frame, we next tested the effect of the inhibitors on cell viability. This variable was examined in the presence and absence of exogenously added glutamine; this is because most cancer cell lines depend on glutaminolysis supporting oxidative decarboxylation and catabolism of glutamine through the citric acid cycle and/or reductive carboxylation towards fatty acid synthesis ^45^. As shown in figure panel 6A, no RC inhibitor increased the death rate of HepG2 cells, irrespective of glutamine availability. *In lieu* of chemical anoxia conferred by azide, we also tested the effect of true hypoxia by subjecting the cultures in oxygen-free environments for 24-72 hours. Mindful of the technical difficulty in achieving true severe hypoxia, we performed immunohistochemical analysis of EF5 ((2-(2-Nitro-1H-imidazol-1-yl)-N-(2,2,3,3,3-pentafluoropropyl) acetamide) adduct formation ^33^. These adducts only form in severely hypoxic cells and can be visualized by applying a fluorophore-conjugated antibody directed against them. As shown in supplementary figure 4, both HepG2 and MCF-7 cultures subjected to 24-72 hours-long hypoxia led to strong labelling by the Alexa 488-conjugated antibody directed against EF5 adducts. Inclusion of EF5 in the absence of hypoxia led to virtually no labelling. Although we cannot accurately estimate the extent of hypoxia, the fact that EF5 adducts appeared to such a degree implies that oxygen concentration must have been <0.1%. Similarly to the effect of azide, hypoxia treatment did not lead to any increase in HepG2 cell death, at least over the course of the experimental time frame (figure panel 6A).

To verify that our method of checking for cell viability (PI/Hoechst33342) is valid, we tested the effects of the pore-forming peptide alamethicin and also that of the detergent digitonin. As shown in figure panel 6B, alamethicin caused a dose-dependent increase in the PI/Hoechst33342 ratio implying loss of plasma membrane integrity which commits a cell to death; digitonin also led to a nearly 100% loss of plasma membrane integrity within 15 minutes of treatment. RC inhibitors, alamethicin or digitonin did not exert a significant effect on total cell number, depicted in figure panels 6C and D. This affords the assurance that our data are not significantly altered by cell detachment; this is important since the method for testing cell viability was high-content automated microscopy, i.e. cells are imaged at specified time intervals.

On the other hand, MCF-7 cells exhibited considerable sensitivity to RC inhibitors, hypoxia or uncoupling; indeed, as shown in figure panel 7A, the presence of rotenone (but not piericidin A not pyridaben), myxothiazol, azide or oligomycin blocking CI, CIII, CIV and CV, respectively, led to a significant elevation in cell death; in a similar manner, true hypoxia or uncoupling by SF6847 also led to a universal cell death of the cultures. To verify that the technique of assessing cell viability is also valid for MCF-7, we tested the effects of alamethicin and digitonin, just as for HepG2 cells. As shown in figure panel 7B, alamethicin caused a dose-dependent increase in cell death; digitonin led to a nearly 100% loss of cell viability. Total cell number of MCF-7 remained unaffected except in the case of digitonin, that led to a drop to ∼25%; this is probably because digitonin detaches MCF-7 cells much more readily than other cell types from the bottom of the plate.

Mindful that aspartate availability has been reported to rescue respiration-deficient cells ^46, 47, 48, 49^, we tested the effect of including an excess of aspartate (10 mM) in the media feeding MCF-7 cells, and check whether this would prevent RC-inhibitor cell death. As shown in figure 8, the inclusion of 10 mM aspartate prevented CII-, CIII- and CIV-inhibitor mediated cell death by up to 15%. Although this is statistically significant, we conclude that aspartate availability cannot be the sole factor in regulating RC inhibitor-mediated MCF-7 cell death.

**Figure 5:**
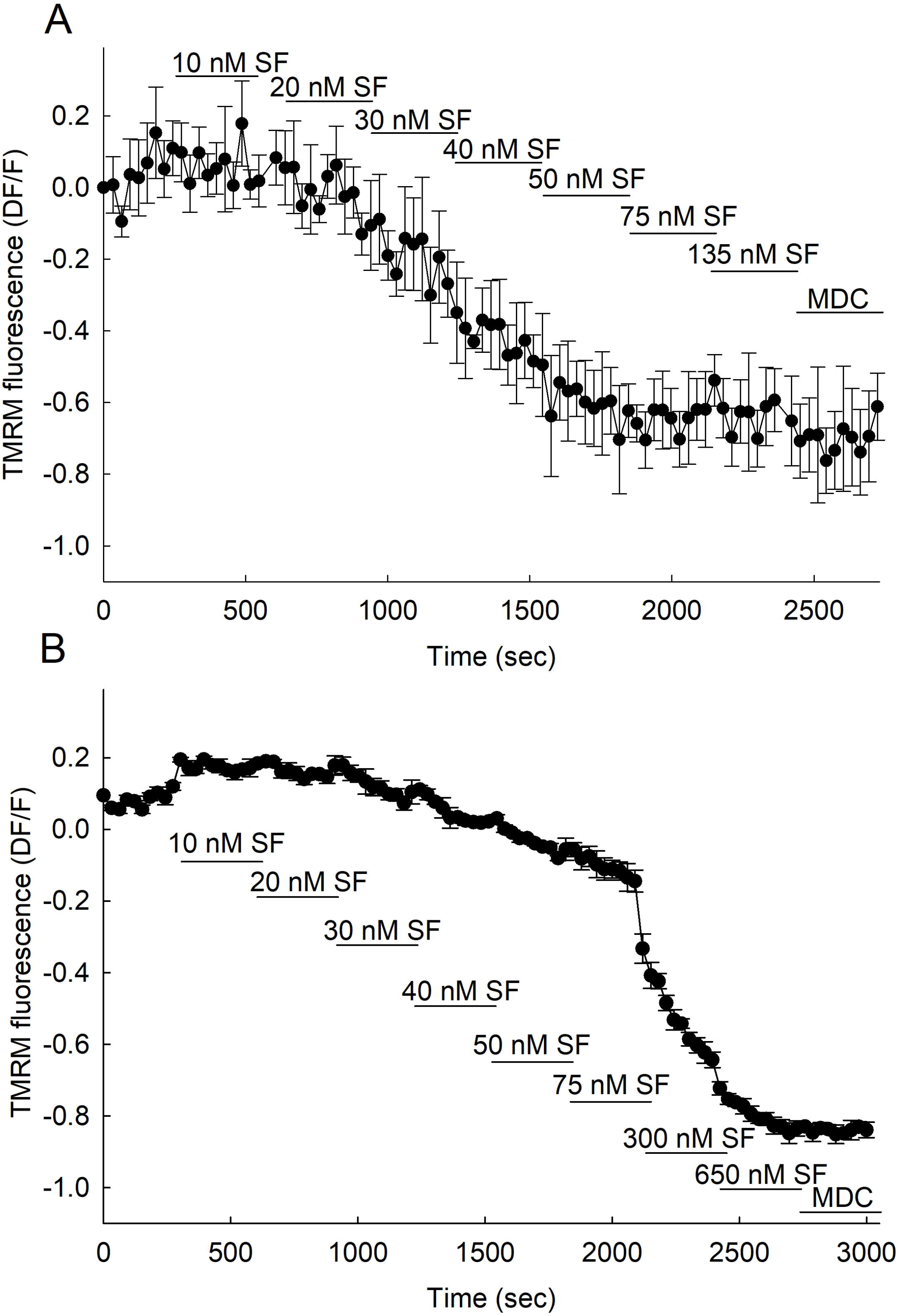
Titration of the uncoupler SF6847 as a function of *in situ* mitochondrial membrane potential (ΔΨm) in HepG2 (A) and MCF7 cells (B). Y axis reflects ΔΨm expressed as the mitochondrial intensity of TMRM fluorescence decomposed of its plasma membrane content. At the end of each experiment full mitochondrial depolarization was achieved by the application of mitochondrial depolarization cocktail (MDC) containing (in μM): 1 valinomycin, 1 SF 6847, 2 oligomycin. Data indicate mean of three independent experiments and error bars indicate SEM from a total of 724 HepG2 and 1561 MCF7 cells.

**Figure 6:**
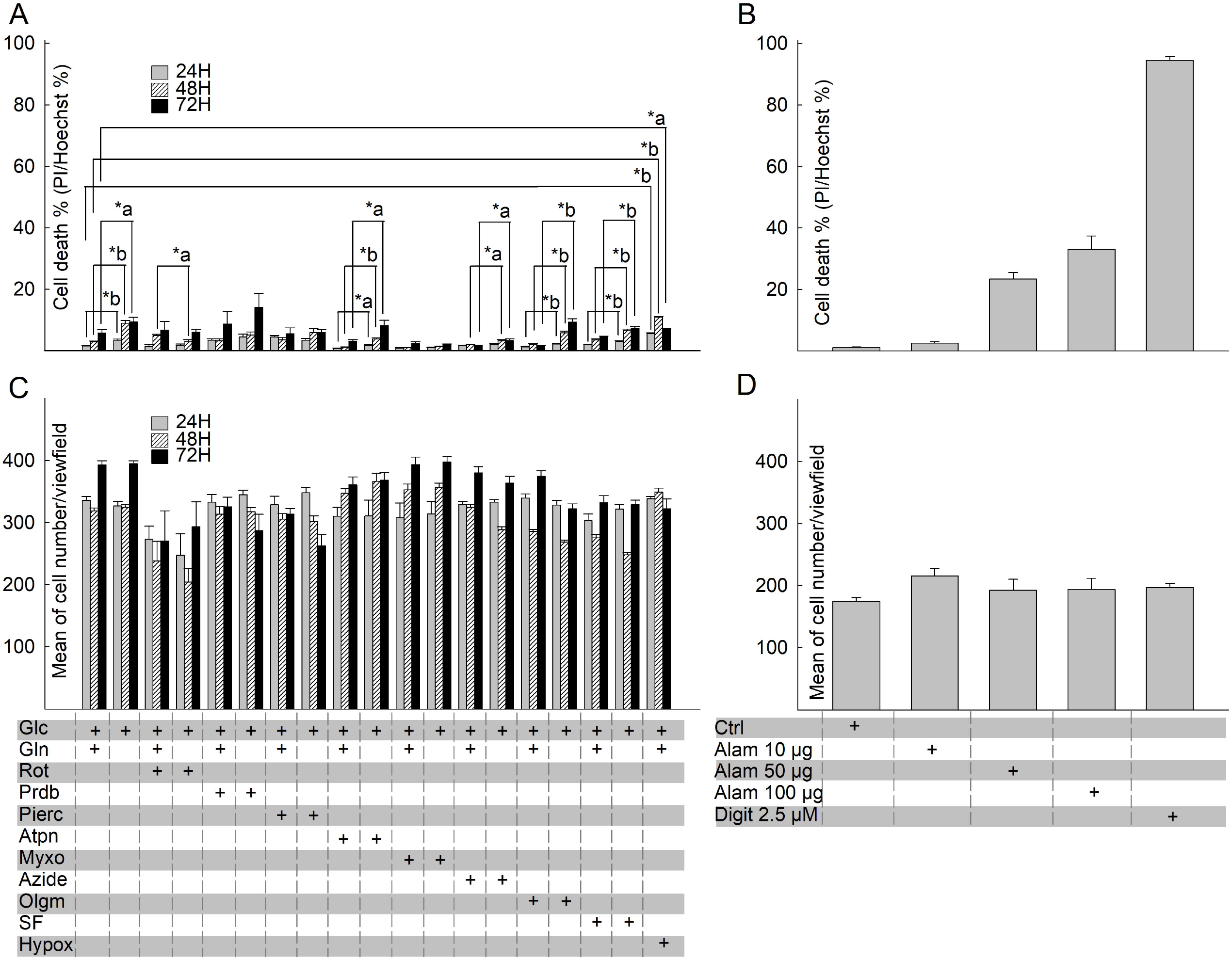
Effect of site-specific RC inhibition on the viability of HepG2 cancer cell line. (**A**) 24h, 48h and 72h site-specific RC inhibition-induced cytotoxicity of HepG2 cells was assessed using nucleus-based quantification of dead (propidium iodide (PI) positive)/ total (Hoechst 33342 (Hoechst) positive) cells. (**B**) Effects of the pore-forming peptide alamethicin (Alam) and the detergent digitonin (Digit) on the viability of HepG2 cells (**C**) Mean number of HepG2 cells/field of view after 24h, 48h and 72h site-specific RC inhibition. (**D**) Mean number of HepG2 cells/field of view after 15 min alamethicin or digitonin application. Bars indicate mean of at least three independent experiments, each addition with 12-32 statistical replicates and error bars indicate SEM. Concentrations of RC inhibitors were identical to those used in the metabolic profiling of HepG2 cells and see it in details in the legend of Fig. 3. *a: p <0.05; *b: p <0.001.

**Figure 7:**
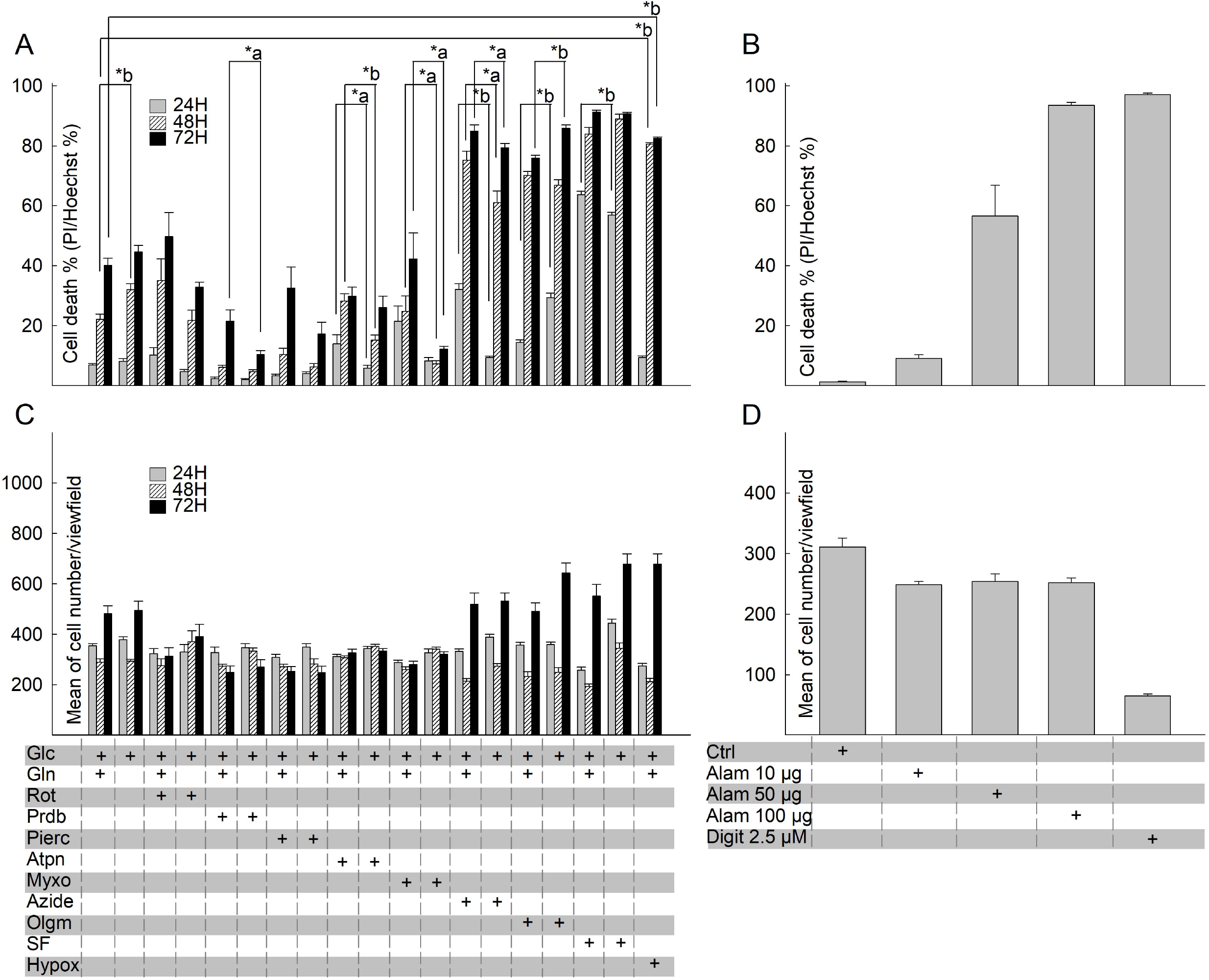
Effect of site-specific RC inhibition on the viability of MCF7 cancer cell line. (**A**) 24h, 48h and 72h site-specific RC inhibition-induced cytotoxicity of MCF7 cells was assessed using nucleus-based quantification of dead (propidium iodide (PI) positive)/ total (Hoechst 33342 (Hoechst) positive) cells. (**B**) Effects of the pore-forming peptide alamethicin (Alam) and the detergent digitonin (Digit) on the viability of MCF7 cells (**C**) Mean number of MCF7 cells/field of view after 24h, 48h and 72h site-specific RC inhibition. (**D**) Mean number of MCF7 cells/field of view after 15 min alamethicin or digitonin application. Bars indicate mean of at least three independent experiments, each addition with 12-32 statistical replicates and error bars indicate SEM. Concentrations of RC inhibitors were identical to those used in the metabolic profiling of MCF7 cells described in the legend of Fig. 3. *a: p <0.05; *b: p <0.001.

**Figure 8:**
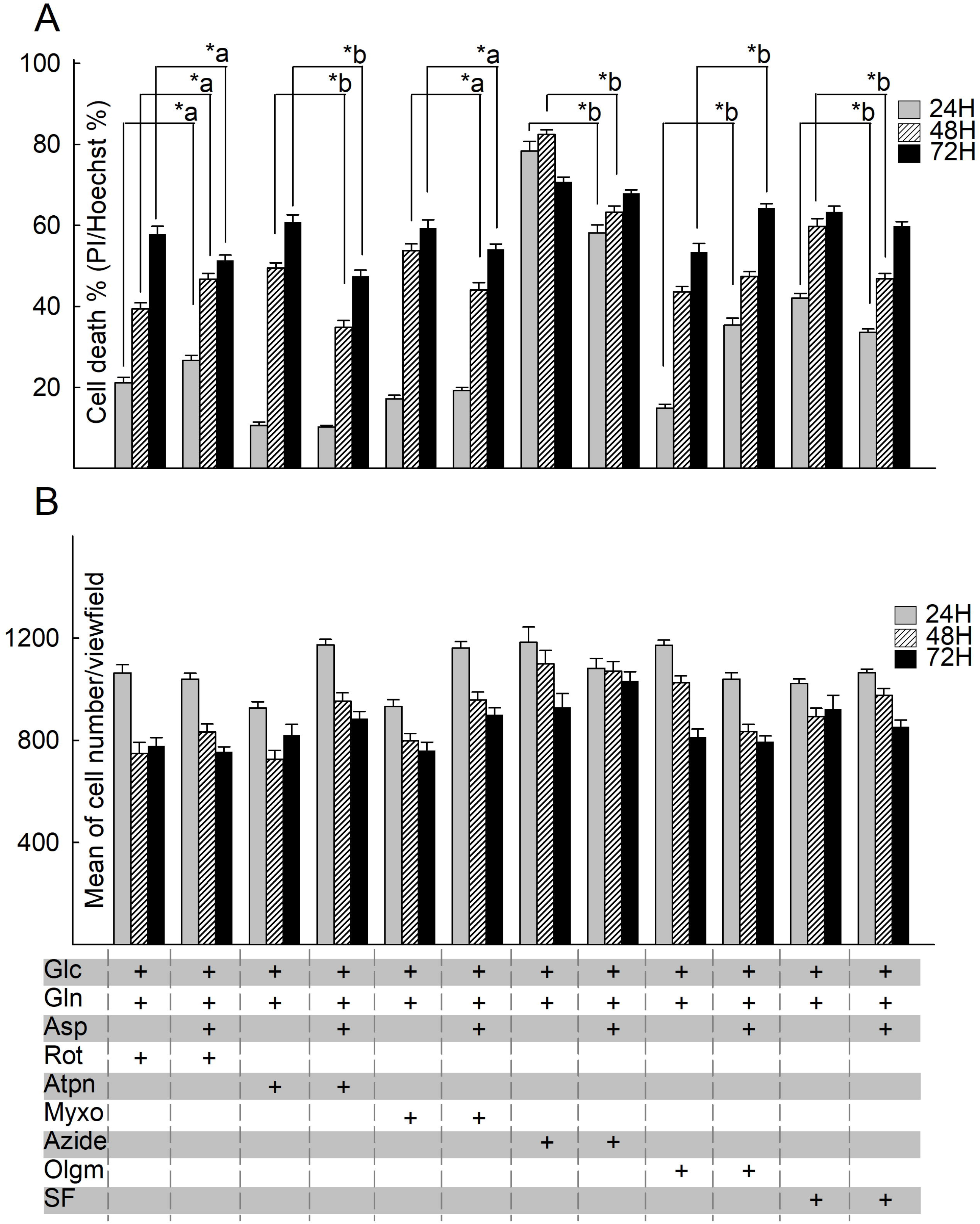
Aspartate supplementation partially restores viability of RC inhibited MCF7 cancer cells. (**A**) 24h, 48h and 72h site-specific RC inhibition-induced cytotoxicity of aspartate supplemented MCF7 cells was assessed using nucleus-based quantification of dead (propidium iodide (PI) positive)/ total (Hoechst 33342 (Hoechst) positive) cells. (**B**) Mean number of MCF7 cells/field of view after 24h, 48h and 72h site-specific RC inhibition. Bars indicate mean of at least three independent experiments, each addition with 12-32 statistical replicates and error bars indicate SEM. *a: p <0.05; *b: p <0.001. Concentrations of RC inhibitors were identical to those used in the metabolic profiling of MCF7 cells and see it in details in the legend of Fig. 3.

## Discussion

The main question addressed in the present work was the impact of bioenergetic impairment on cancer cells viability. To this end, the nature of the bioenergetic impairment was exhaustively addressed by using site-specific poisons of mitochondrial functions. Hereby, drugs were used that would either impair the ability of mitochondria to i) develop membrane potential (using an uncoupler) or ii) maintain a protonmotive force (using CI-CIV inhibitors or severe hypoxia), or iii) perform ATP synthesis (using a CV inhibitor). Each and every regiment aimed to prevent mitochondria from yielding ATP by OXPHOS: SF6847 would uncouple electron transport from ATP synthesis and also abolish the ATP-yielding directionality of the F_o_-F_1_ ATP synthase, the inhibitors CI-CIV or severe hypoxia would inhibit the electron transport thus abolish the protonmotive force, while oligomycin would prevent ATP formation directly by inhibiting the F_o_-F_1_ ATP synthase. Our results unequivocally show that no bioenergetic impairment led to a decrease in viability of HepG2 cells, while these cultures exhibited strong OXPHOS activity and respiratory capacity. On the other hand, MCF-7 cultured cells viability exhibited sensitivity to some bioenergetic impairment regiments, but also showed very weak OXPHOS. This lack of correlation urges for caution when linking cancer cell survival as a function of OXPHOS, also arguing that no extrapolations can be made from one cell type or condition, to another. Relevant to this, it has been recently reported that the tumor-growth inhibitory activity of complex I poisons is not due to energy depletion; instead they mitigate cancer growth by altering the pH of the tumor microenvironment ^50^. The following findings deserve further scrutiny:

i. why would HepG2 cells invest in expressing all respiratory chain components, be capable of OCR, ECAR, ΔΨm and yet be resistant to any bioenergetic impairment regime that would abolish mitochondrial ATP efflux? In the literature, the list of moonlighting functions by mitochondrial proteins is constantly expanding ^51, 52, 53^, however, it is puzzling (at least to these authors) as to why express the components of the entire respiratory chain, be functional, and yet not need it.
ii. by the same token, why would MCF-7 cells exhibit so little -if any-mitochondrial bioenergetic competence to the point that it is doubtful that they need OXPHOS, and yet be so sensitive to some (but not all) bioenergetic impairment regimes? To this end, it may be possible that MCF-7 are critically dependent on this meagre bioenergetic OXPHOS, and by abolishing it -no matter how little-is detrimental to them. Relevant to this, the effect of severe hypoxia or CIV inhibition but not CIII in conferring MCF-7 cell death is puzzling; in theory, CIV could be fuelled by metabolites such as glutathione or proteins (such as p66Shc), when CIII is inhibited ^54, 55^. Such possibilities may be worth investigating.
iii. the residual OCR in MCF-7 cells after the acute addition of antimycin and rotenone blocking CIII and CI respectively, must be highlighted. This points to a non-mitochondrial component consuming oxygen; thus, whenever oxygen consumption is observed, it should not automatically be attributed to mitochondria, let alone OXPHOS.

Overall, our data highlight that the relation between OXPHOS and cell viability is not straightforward, and it must be examined in cell- and condition-specific manner. The importance of this is exemplified by the evolving concept that cancer could be managed by metabolic drugs ^44^.

## Acknowledgements

This work was supported by grants from NKFIH (KH129567, and K135027) to C.C. We thank the Department of Physiology of Semmelweis University for providing access to the ImageXpress Micro Confocal High Content Imaging System (supported by VEKOP-2.3.2-16-2016-00002).

