## Supplementary material for "Viability of HepG2 and MCF-7 Cells is not Correlated with Mitochondrial Bioenergetics": legends to suppl figures

*Supplementary figure legends:*

**Supplementary Figure 1: Membrane polarisation loss with increasing azide concentrations in anoxia.** A representative trace of membrane potential determination (**A**) with isolated mouse mitochondria. Additions and the depletion of oxygen (anox) are indicated on top: 2 mM ADP, varying concentrations of azide, 250 nM SF 6847 (uncoupler) respectively. Blue trace represents the oxygen concentration; red trace represents the rhodamine 123 fluorescence indicative of membrane potential. Simultaneous plot of the % anoxic membrane potential loss (red) and % remaining cytochrome c oxidase catalytic activity (green) (**B**). *Fl*_anox_, *Fl*_azide_ and *Fl*_unc_ are fluorescence values at the end of anoxic, azide inhibited and uncoupled state respectively. Values greater than 100% mean all the polarisation was lost, but there is a gradual decay in fluorescence over time as can be seen on (**A**) after the uncoupler*. k’*_0 mM_, *k’*_5 mM_ and *k’_c_* are the pseudo first order rate constants at 0 mM, 5 mM and varying concentrations of azide respectively, measured in HepG2 cells. Data points indicate median of 3-5 measurements and error bars indicate 1 standard deviation around the mean.

**Supplementary Figure 2: Effect of combined inhibition of CI+ CV or CIII+CV on *in situ* mitochondrial membrane potential (ΔΨm) in HepG2 (A, B) and MCF7 cells (C,D)**. Y axis reflects ΔΨm expressed as the mitochondrial intensity of TMRM fluorescence decomposed of its plasma membrane content. Addition of each inhibitor is indicated in the graph and at the end of each experiment full mitochondrial depolarization was achieved by the application of mitochondrial depolarization cocktail (MDC) containing: 1 μM valinomycin, 1 μM SF 6847, 2 μM oligomycin. Data indicate mean of three independent experiments and error bars indicate SEM.

**Supplementary Figure 3: Citrate synthase catalytic activity content of MCF7 cells over time.** Citrate synthase measured after 24h, 48h or 72h at 37 °C under 5% CO_2_ atmosphere in assay media. Inhibitor treated (grey) and their respective control (black) cells were normalized to protein equivalent. Inhibitors were 5 μM rotenone (**A**), 1 μM atpenin A5 (**B**), 1 μM myxothiazol (**C**), 1 mM azide (**D**), 5 μM oligomycin (**E**)**.** Bars indicate the mean of three biological replicates and error bars indicate 1 standard deviation. *a: p <0.05; *b: p <0.001.

**Supplementary Figure 4: Effect of 24h, 48h and 72h of hypoxia on a hypoxia marker (EF5 adduct) formation in HepG2 and MCF7 cells.** EF adducts (EF add.) stain appears only during hypoxia. Phase contrast (phase) images verify the presence of cells when there is no EFF add. staining.
