## Supplementary figures and images for "Viability of HepG2 and MCF-7 Cells is not Correlated with Mitochondrial Bioenergetics"

### Supplementary figure 1

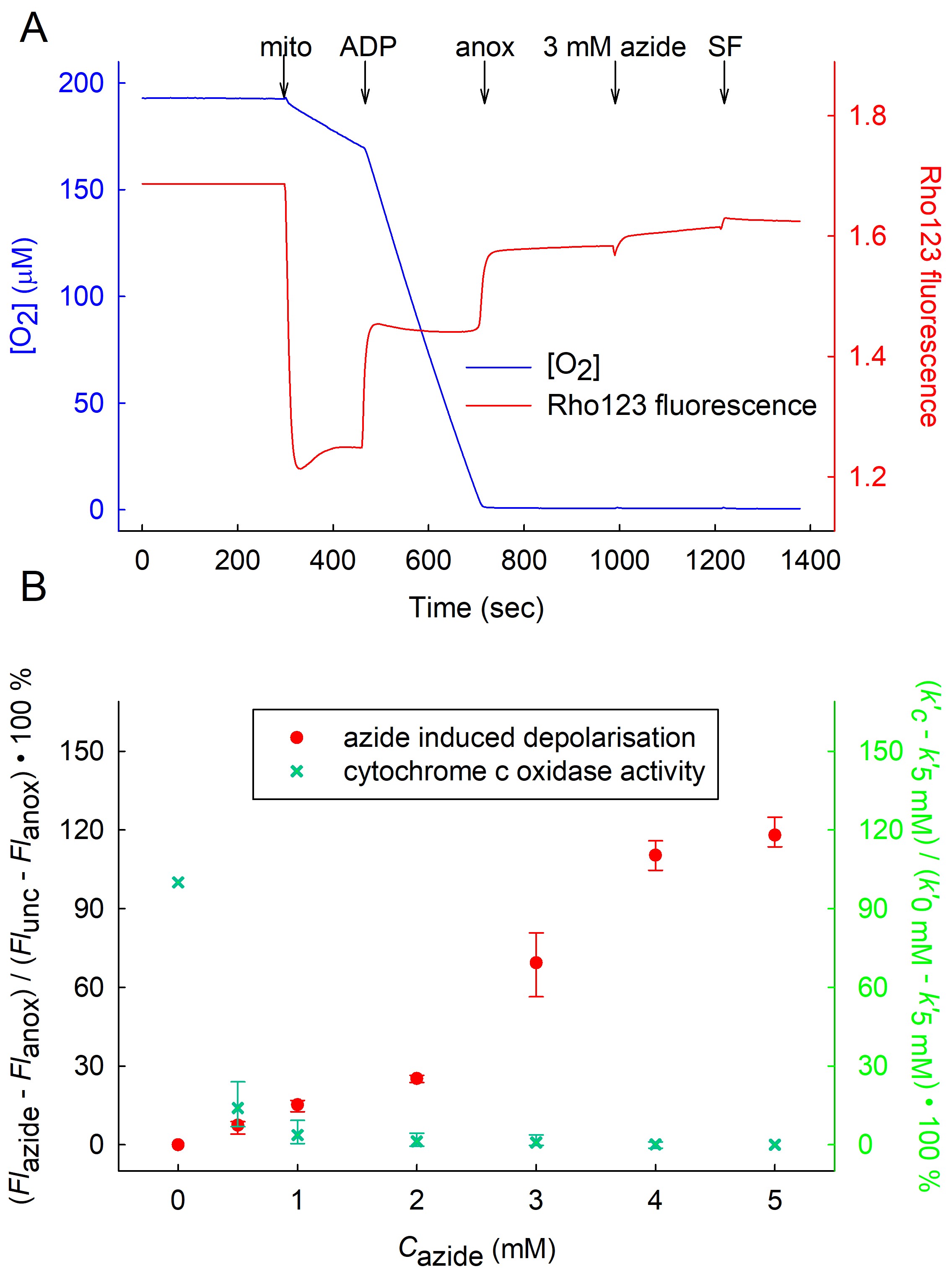

### Supplementary figure 2

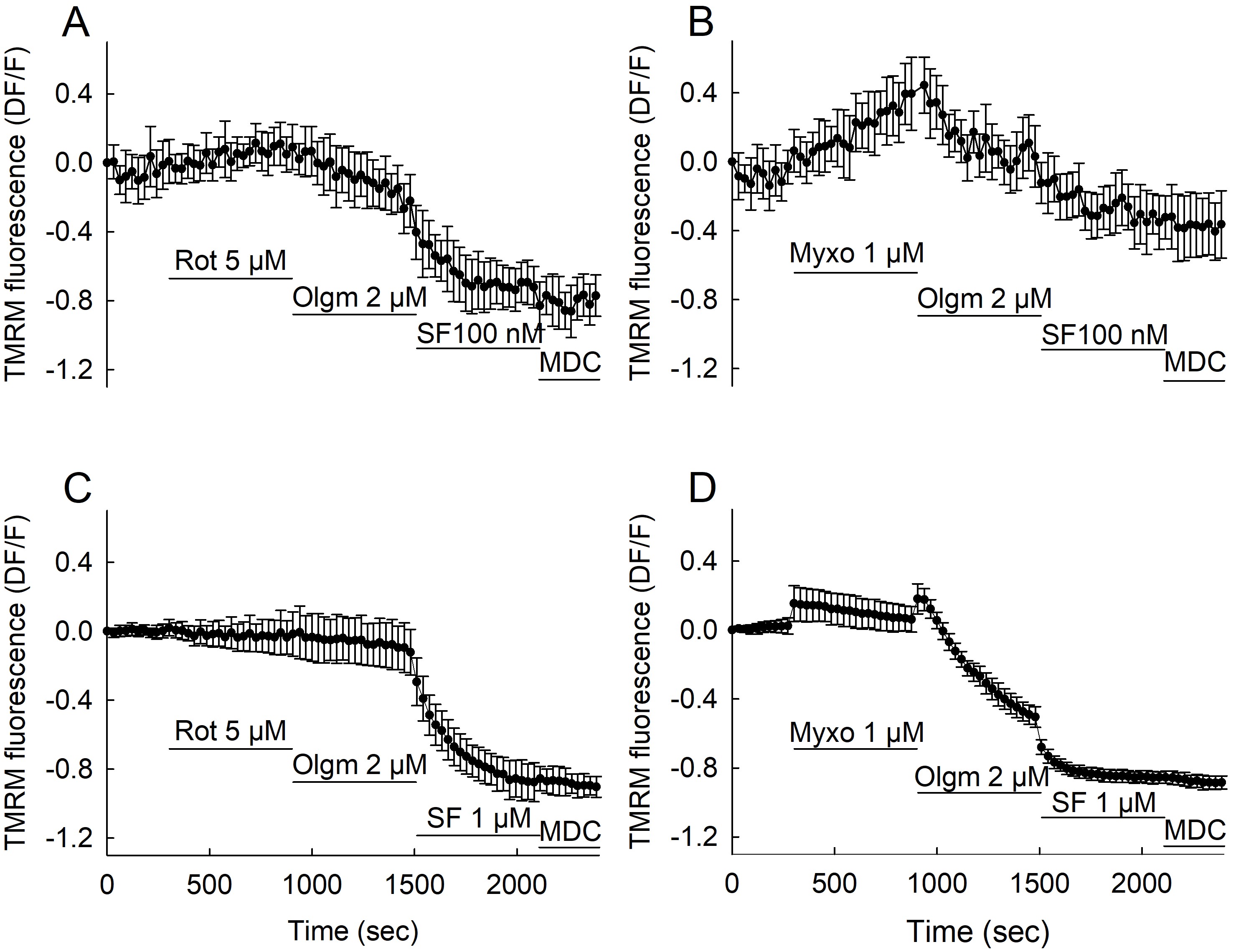

### Supplementary figure 3

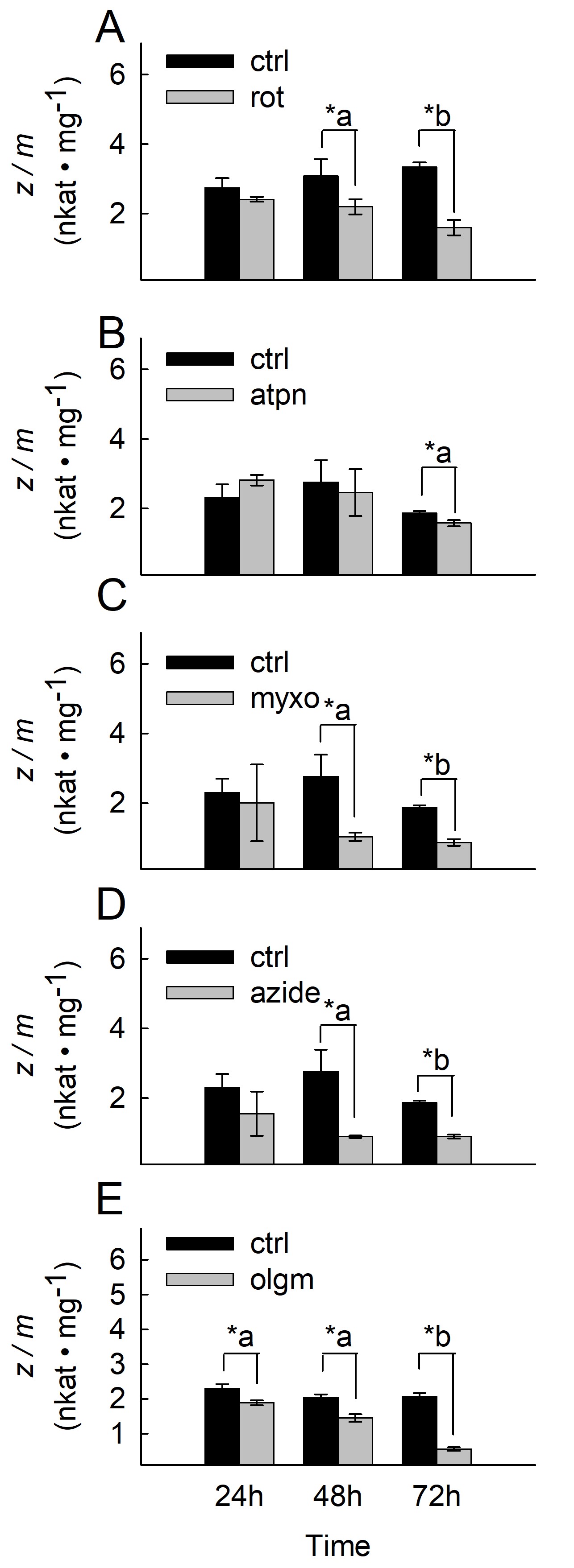

### Supplementary figure 4

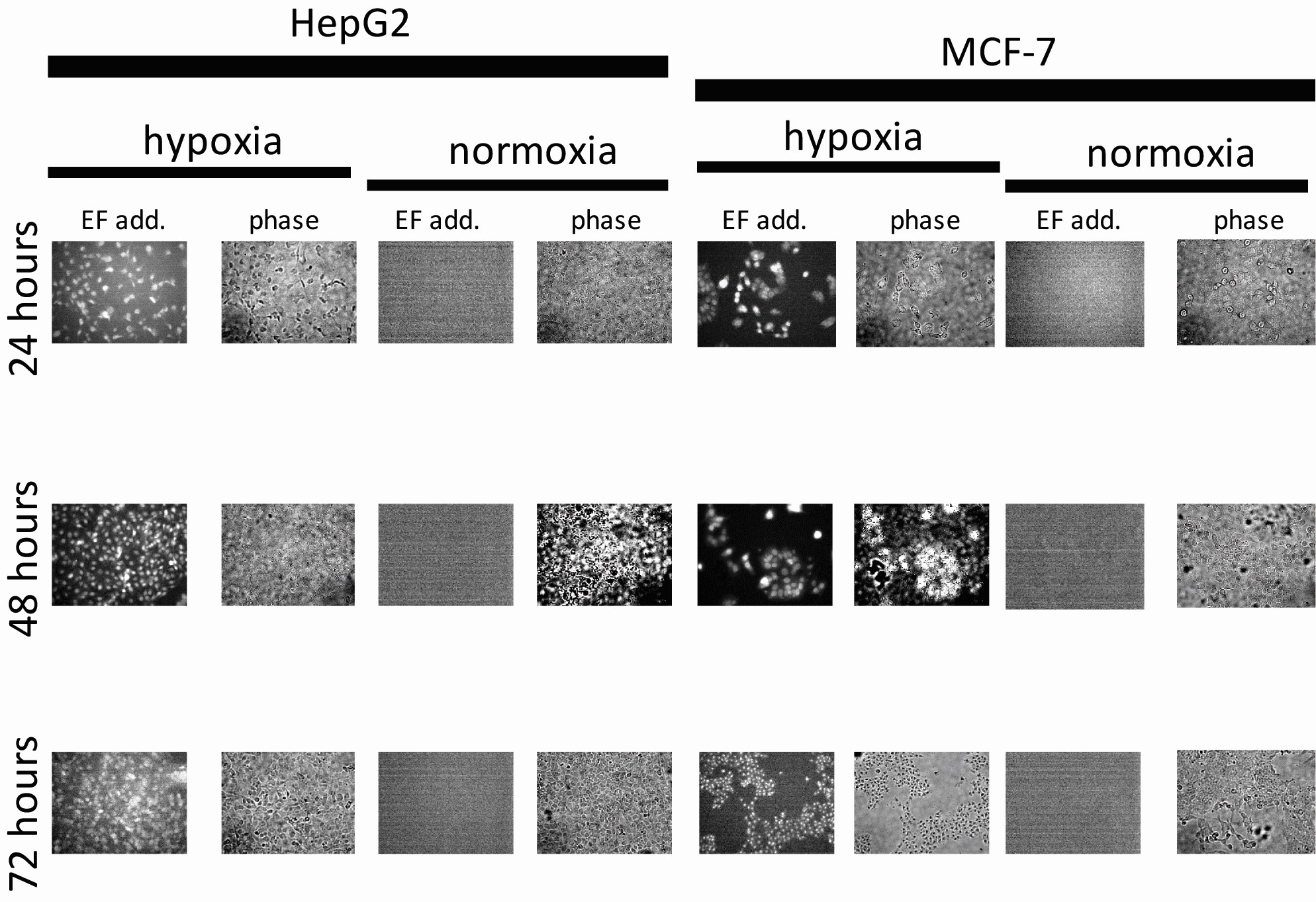
